# EML4–ALK Fusion Rewires Transcriptomic, miRNA, and CAF-Associated Programs in Non-Small Cell Lung Cancer

**DOI:** 10.64898/2026.04.10.717853

**Authors:** Divya Mishra, Shivangi Agrawal, Divya Malik, Ekta Pathak, Rajeev Mishra

## Abstract

Graphical abstract

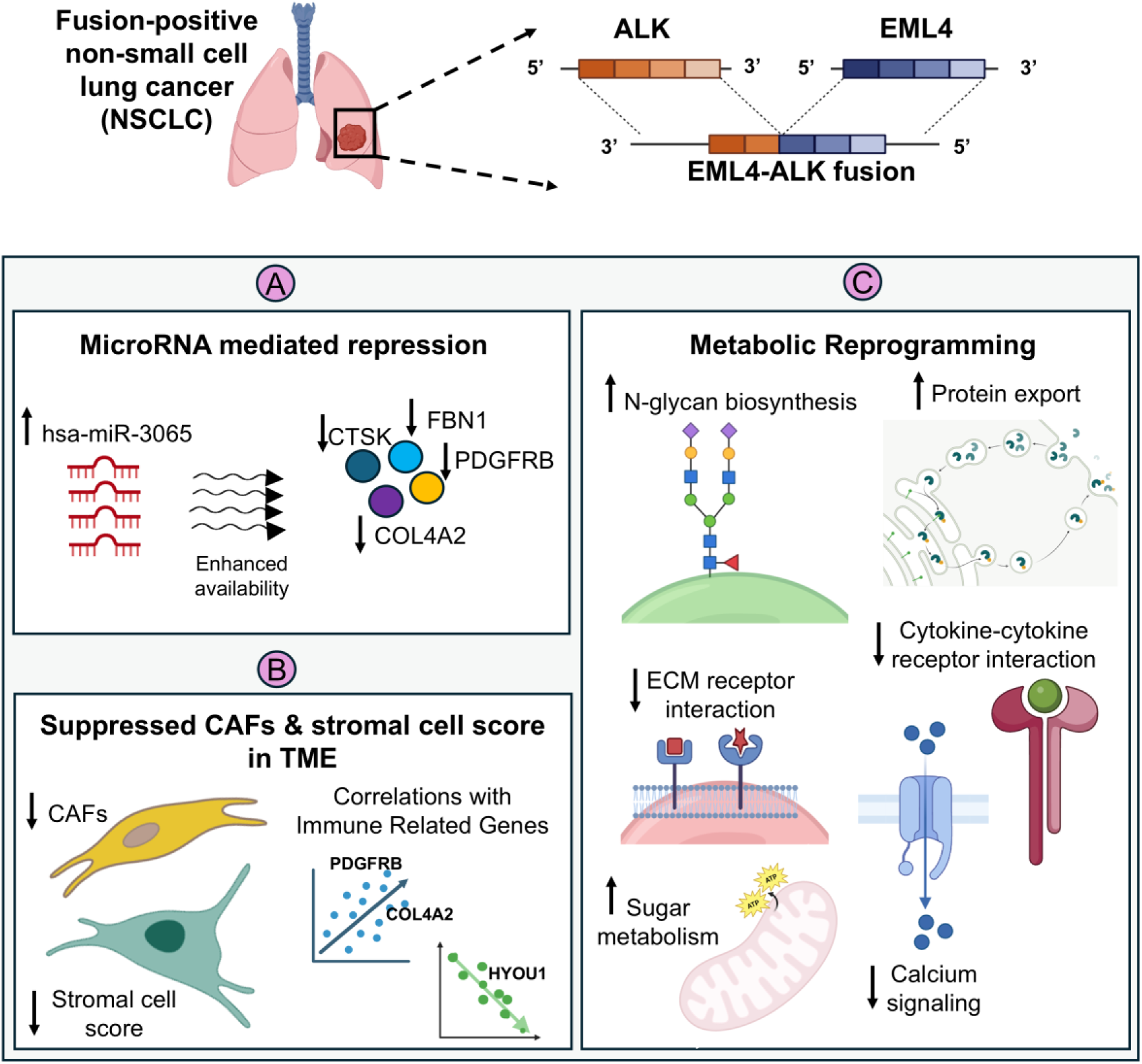

This study establishes an integrative framework that combines paired mRNA/miRNA profiling with immune microenvironmental features to clarify how EML4–ALK fusions shape transcriptomic and post-transcriptional networks in Non-small cell lung cancer (NSCLC). Using paired mRNA-seq and miRNA-seq data generated from the same patients, we compared fusion-positive and fusion-negative NSCLC across three interconnected layers: (i) transcriptome architecture, including differential expression, pathway, and network analyses; (ii) the miRNA–mRNA regulatory axis, encompassing dysregulated miRNAs, target repression and sponging, and fusion-specific regulatory pairs; and (iii) the tumor microenvironment, with emphasis on immune and stromal infiltration, particularly cancer-associated fibroblast (CAF)-linked extracellular matrix (ECM) and adhesion programs. Our analyses revealed a distinct reprogramming pattern in fusion-positive NSCLC, marked by activation of metabolic and proteostasis pathways, including N-glycan metabolism coupled to ER export, together with attenuation of immune–stromal communication, adhesion, ECM, calcium signaling, and PI3K/VEGF-axis transcription relative to fusion-negative NSCLC. We also identified fusion-associated microRNA perturbations, including an exclusively upregulated miR-3065-centered regulatory hub predicted to repress ECM- and adhesion-related genes (PDGFRB, CTSK, COL4A2, SPARC, FBN1, and LUM) in fusion-positive tumors, in contrast to broader miRNA network rewiring in fusion-negative tumors targeting ciliary and mitotic hubs. Tumor microenvironment analysis further distinguished the subtypes, with fusion-positive tumors showing reduced CAF infiltration relative to fusion-negative tumors and concordant gene–CAF associations. By linking mechanistic insight with candidate biomarkers and targetable pathway nodes, this work provides a basis for precision strategies in both fusion-positive and fusion-negative cohorts and broadens the therapeutic perspective beyond kinase inhibition alone.

## Introduction

Non-small cell lung cancer (NSCLC) accounts for ∼85% of lung cancers and remains the leading cause of cancer mortality. Among the key oncogenic drivers of NSCLC is the EML4-ALK fusion, present in ∼4-7% of NSCLCs, particularly lung adenocarcinomas, generated by a chromosomal inversion on 2p that joins the EML4 N-terminus to the ALK kinase domain^1–3^. This rearrangement produces a constitutively active kinase that engages major growth and survival pathways, including RAS/RAF/MEK/ERK and PI3K/AKT/mTOR, defining a molecular subtype that is overrepresented in younger patients and never or light smokers and is largely mutually exclusive with EGFR or KRAS alterations ^4^. Although ALK tyrosine-kinase inhibitors (TKIs) have substantially improved clinical outcomes, clinical heterogeneity and acquired resistance highlight that protein-centric signaling models incompletely explain fusion-driven biology ^5^.

Accumulating evidence suggests that oncogenic fusions also remodel RNA-level regulation, reshaping the tumor transcriptome beyond kinase signaling ^6–11^. A central component of this regulatory layer is microRNAs (miRNAs), ∼22-nt molecules that repress target transcripts through 3′-UTRs ^12–15^. In cancer, aberrant miRNA expression can confer oncogenic or tumor-suppressive functions, broadly influencing cell proliferation, apoptosis, metastasis, and immune evasion ^16,17^. In fusion-driven malignancies, changes in miRNA abundance and target availability can propagate transcriptome-wide effects. At a mechanistic level, fusion transcripts can reshape 3′-UTR architecture, simultaneously eliminating native miRNA binding sites and, when highly expressed, redistributing miRNA binding and sponging effects, thereby amplifying dysregulation at the network level ^18–21^. In the context of EML4-ALK, loss of the EML4 3′-UTR together with elevated ALK transcript expression may perturb miRNA target landscapes by removing native EML4 binding sites and altering miRNA partitioning toward abundant ALK 3′-UTRs, thereby reinforcing fusion-dependent transcriptional programs. Together, these observations prompt us to study the miRNA-mRNA axis as a critical determinant of how ALK fusions sustain malignant programs beyond kinase output.

The tumor immune microenvironment (TIME) adds an additional layer of complexity ^22^. NSCLC tumors comprise complex ecosystems of diverse immune lineages, including T, B, and NK cells, macrophages, and neutrophils, alongside stromal components such as endothelium and cancer-associated fibroblasts (CAFs), that influence tumor progression and responses to therapy ^23,24^. MicroRNAs also play key roles in shaping stromal phenotypes and mediating cancer-stroma crosstalk ^25–27^. How EML4-ALK fusions integrate miRNA regulation with immune microenvironmental states to shape clinical behavior has not yet been fully elucidated.

Here, we performed a fusion-stratified integrative analysis of paired mRNA-seq and miRNA-seq data from the same NSCLC patients to define how EML4–ALK fusion status shapes transcriptomic, post-transcriptional, and tumor microenvironmental states. We compared fusion-positive and fusion-negative tumors across three interconnected layers: global gene-expression architecture, the miRNA–mRNA regulatory axis, and immune–stromal composition, with particular attention to cancer-associated fibroblast- and extracellular matrix-related programs. This framework was used to identify fusion-associated pathways, regulatory hubs, and candidate biomarkers that may help explain the biological divergence between these molecular subtypes and support more precise therapeutic stratification.

## Materials and Methods

### Data acquisition and processing

RNA sequencing (RNA-seq) and microRNA sequencing (miRNA-seq) profiles for non-small cell lung cancer (NSCLC), were obtained from The Cancer Genome Atlas (TCGA) (https://www.cancer.gov/tcga). Tumor and adjacent normal tissue data were downloaded through the Genomic Data Commons (GDC) portal (https://portal.gdc.cancer.gov). Specifically, STAR-Counts files were used for RNA-seq data, which included both protein-coding and long non-coding RNAs, while miRNA expression was derived from British Columbia Genome Sciences Centre (BCGSC) profiling files. Samples harboring the EML4-ALK fusion were identified with the ChimerSeq module of ChimerDB 4.0 ^28^. Based on the fusion status, patients were stratified into EML4-ALK fusion-positive or fusion-negative groups. In total, RNA-seq data were available for 474 lung adenocarcinoma and adenoma samples and 59 matched normal tissues. Of these, five tumor samples were fusion-positive and 469 were fusion-negative. For downstream analyses, we focused on 66 non-smoker, fusion-negative tumors as a comparator group. Differential expression was evaluated across two contrasts: (i) fusion-positive NSCLC vs. normal and (ii) fusion-negative NSCLC vs. normal.

For miRNA-seq, five fusion-positive tumors, 61 non-smoker fusion-negative tumors, and 46 normal samples were available. Only samples with both RNA-seq and miRNA-seq data were included for integrative analyses. Quality control involved filtering out low-count genes and applying principal component analysis (PCA) and multidimensional scaling (MDS) to detect outliers and visualize cohort structure. Outlier analysis was conducted using factoextra and the plotMDS function in LIMMA^29,30^. After these steps, four fusion-positive and 61 fusion-negative NSCLC samples with paired datasets remained.

### Differential gene and miRNA expression analyses

DESeq2 employed Wald tests, whereas LIMMA normalized raw counts into log2 counts per million (logCPM) and applied a linear modeling framework. Genes were considered differentially expressed when |log2 fold change| ≥ 1 and padj < 0.05. Final DEG lists were generated by intersecting results from both tools. Similarly, differentially expressed miRNAs (DEMs) were identified under the same thresholds. Exclusive expression signatures for fusion-positive and fusion-negative groups were derived by comparative analyses. Volcano plots were generated with EnhancedVolcano ^31^, and overlaps visualized with InteractiVenn ^32^.

### Protein-protein Interaction (ppi) network

PPI networks were generated separately for upregulated and downregulated DEGs in both fusion-positive and fusion-negative NSCLC using the GeneMANIA application in Cytoscape ^33,34^. Hub genes were identified with the MCC algorithm in the cytoHubba plugin ^35^. Network modules with high connectivity were extracted using the MCODE algorithm (parameters: degree cutoff = 2, node score cutoff = 0.2, K-core = 2, maximum depth = 100) ^36^. In MCODE, the degree of gene association in the module was scored using the degree cutoff = 2, node score cutoff = 0.2, K-score = 2, and max depth = 100. Hub genes were further examined for overlap with highly connected modules.

### Functional and pathway enrichment analyses

The top three statistically significant gene modules from each PPI network were subjected to functional annotation using DAVID ^37,38^. Enrichment analyses included Gene Ontology (GO) categories-biological process (BP), molecular function (MF), and cellular component (CC)-as well as KEGG pathways. Terms with p < 0.05 were considered significant, and results were visualized as bubble plots with ggplot2 ^39^.

### Gene set enrichment analysis (GSEA)

To further interrogate biological patterns, GSEA was performed using the c2.cp.kegg.v2023.1.Hs.symbols.gmt collection from MSigDB ^40,41^. Analyses contrasted fusion-positive with fusion-negative NSCLC, with 1000 phenotype permutations. All other parameters were kept at default settings.

### Prediction of miRNA targets

MicroRNAs identified from TCGA were first standardized to their mature forms by appending 3p/5p annotations using the miRBase database ^42^. The mature miRNA set was then queried in the miRDiP 4.1 database to predict targets of differentially expressed miRNAs (DEMs) ^43^. Additionally, candidate miRNAs targeting the EML4 and ALK genes, along with their putative downstream targets, were retrieved. Predictions were filtered to retain only 3′ UTR-localized binding sites and limited to pairs supported by at least three databases, using the medium-confidence threshold.

### Identification of differentially expressed targets (DETS)

Predicted miRNA-gene interactions were cross-referenced with differentially expressed genes (DEGs) to define differentially expressed targets (DETs). Exclusive DETs for fusion-positive NSCLC were obtained through comparative Venn diagram analysis.

### Construction of miRNA-mRNA regulatory networks

Regulatory networks linking upregulated DEMs to downregulated targets, and downregulated DEMs to upregulated targets, were assembled and visualized using Cytoscape v3.10.1 ^33,34^. Hub DEMs were identified by degree centrality, and connections between hub DEMs and hub DEGs from the PPI network were highlighted. Interactions involving dysregulated miRNAs targeting EML4 or ALK were also extracted.

### Computational assessment of miRNA-mRNA interactions

Mature miRNA sequences were retrieved from miRBase, and 3′ UTR sequences of target genes downloaded in FASTA format from Ensembl (https://asia.ensembl.org). Binding interactions were evaluated in silico using RNAhybrid ^44^ with miRNA-mRNA pairs retained at mfe ≤ −18 kcal/mol. Interaction strengths were visualized as heatmaps with the pheatmap package in R ^45^.

### Structural modeling of miRNA-AGO2 complexes

Secondary structures of miRNA-mRNA duplexes were predicted with RNAhybrid ^44^. The sequence of human Argonaute 2 (HsAGO2) was retrieved from UniProt ^46^, given its role in guiding miRNAs for transcript cleavage ^47^. Ternary complexes of miRNA-mRNA duplexes bound to HsAGO2 were modeled with AlphaFold 3 ^48^. Predicted complexes were ranked by interface predicted template modeling (ipTM) and predicted template modeling (pTM) scores, applying thresholds of ipTM ≥ 0.8 and pTM ≥ 0.9. The highest-confidence models were further analyzed and visualized using UCSF Chimera v1.18 ^49^.

### Estimation of tumor purity and immune/stromal infiltration

Tumor purity, stromal content, and immune infiltration were quantified in fusion-positive and fusion-negative NSCLC samples using the ESTIMATE R package ^50^. Stromal and immune scores were derived from normalized gene expression matrices, with the combined output representing the ESTIMATE score. In addition, infiltration levels of immune and stromal populations-including total T cells, CD8⁺ T cells, NK cells, B cells, macrophages, myeloid dendritic cells, neutrophils, endothelial cells, and cancer-associated fibroblasts-were estimated using the MCP-counter deconvolution method implemented in the Immunedeconv R package ^51,52^.

### Retrieval of immune-related genes

Immune-related genes were compiled from the Immunome and ImmPort databases ^53,54^, supplemented with CAF-associated gene lists from recent studies. Differential expression analysis was performed to identify immune-related genes altered in EML4-ALK fusion-positive versus fusion-negative NSCLC.

### Correlation analysis of immune infiltration and hub genes

Associations between immune infiltration levels and ESTIMATE-derived scores were assessed using Spearman’s correlation, implemented in the corrplot R package ^55^. Correlations with coefficient > 0.7 and p < 0.05 were considered significant.

### Receiver operating characteristic (ROC) analysis

Genes and miRNAs prioritized from PPI and functional analyses were subjected to ROC analysis. Normalized expression values were obtained using the estimateSizeFactors function in R. ROC curves and AUC values were computed with the pROC package ^56^, and visualized using the plotROC function in ggplot2 ^39^. Features with AUC ≥ 0.7 were considered predictive, while those with overstated values (AUC = 1) were excluded.

### Drug repurposing via connectivity map (CMap)

Candidate small molecules with potential therapeutic activity were identified using the CMap platform ^57^. Module-specific analyses were conducted: for fusion-positive cases, upregulated and downregulated DEGs from modules 1-3 were queried separately; for fusion-negative cases, modules 1 and 2 were merged due to insufficient DEGs, while module 3 was analyzed independently. Each query was run against the Touchstone L1000 reference signatures. Compounds were ranked by normalized connectivity scores, with negative scores reflecting reversal effects. The top 20 negatively scored molecules per query were retained, excluding those with undefined mechanisms of action. Drug targets were extracted, and only molecules whose targets overlapped with significantly upregulated genes or dysregulated pathways in our dataset were prioritized.

## Results

### EML4–ALK fusion status stratifies the NSCLC transcriptome

We analyzed RNA-seq data from EML4–ALK fusion-positive tumors, fusion-negative tumors, and patient-matched normal lung samples. Principal-component analysis clearly separated fusion-positive tumors from normal lung samples (PC1 = 51.0%, PC2 = 10.9%) and similarly distinguished fusion-negative tumors from normal controls (PC1 = 41.1%, PC2 = 12.5%; Figure 1A). Differential expression was assessed using DESeq2 and LIMMA packages. Volcano plots illustrate the top ten up- and down-regulated genes identified by each method (Figure 1B). Intersecting results from both methods identified 3,134 differentially expressed genes (DEGs) in fusion-positive versus normal comparisons (1,238 upregulated; 1,896 downregulated) and 3,451 DEGs in fusion-negative versus normal comparisons (1,788 upregulated; 1,663 downregulated; Figure 1C). Within this set, 569 upregulated and 741 downregulated protein-coding genes were exclusive to fusion-positive NSCLC, whereas 843 upregulated and 455 downregulated genes were exclusiove to fusion-negative cases; 573 upregulated and 1,067 downregulated genes were shared between fusion and non-fusion cohorts (Figure 1C). Hierarchical clustering of the 25 most up- and down-regulated genes per comparison resolved distinct expression programs across sample groups (Figure 1D). Collectively, EML4–ALK fusion status delineates distinct transcriptional programs in NSCLC.

**Figure 1.**
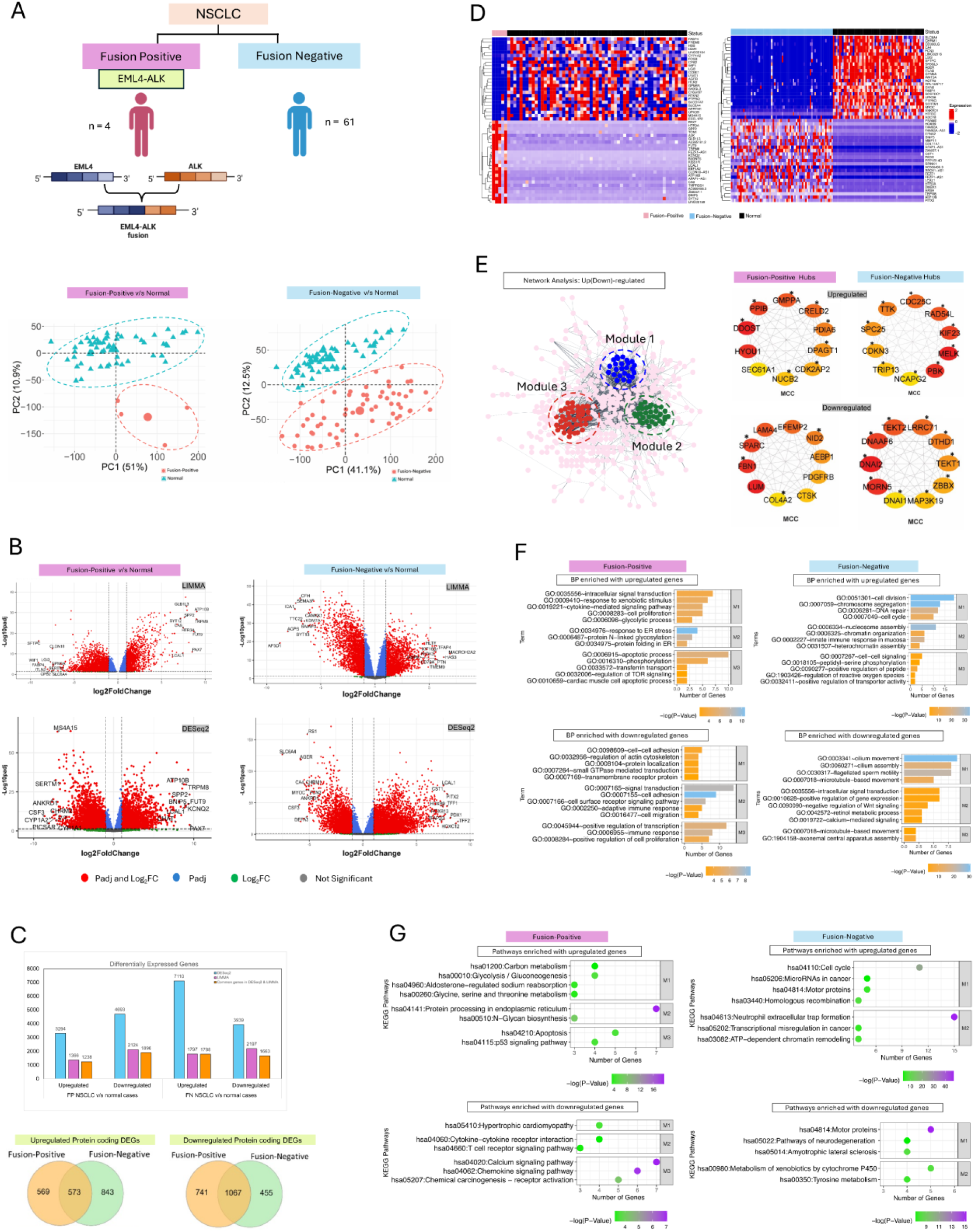
Transcriptomic and network distinctions by EML4-ALK fusion status in NSCLC. (A) Principal-component analysis (PCA) separates fusion-positive versus normal and fusion-negative versus normal samples, with the percent variance explained by PC1/PC2 indicated. (B) Volcano plots for both comparisons showing differentially expressed genes (DEGs) identified by DESeq2 and limma; top up- and down-regulated genes are annotated. (C) Bar chart of DEGs jointly called by both pipelines and Venn diagram indicating subtype-exclusive and shared protein-coding DEGs. (D) Unsupervised hierarchical clustering heat maps of the top 25 up- and down-regulated genes per comparison. (E) Schematic of protein-protein interaction (PPI), representative module maps, and hub-gene marked by Maximal Clique Centrality (MCC). (F) Selected GO Biological Process (GO:BP) terms enriched among up- and down-regulated DEGs for fusion-positive and fusion-negative tumors. (G) KEGG pathway enrichment for up- and down-regulated DEGs in each subtype. See Supplementary Tables S1 and S2 for full gene and pathway lists.

### Interactome topology reveals fusion and non-fusion-specific hub genes

To place these transcriptomic changes in a functional context, we constructed protein–protein interaction (PPI) networks from fusion- and non-fusion-specific DEGs (Figure 1E). In fusion-positive tumors, the resulting networks comprised 569 upregulated and 741 downregulated genes. Three highly connected modules were identified, including upregulated modules containing 103, 15, and 100 nodes, and downregulated modules containing 121, 57, and 76 nodes. Maximal clique centrality (MCC) scoring identified ten upregulated hub genes, including HYOU1, DDOST, PPIB, GMPPA, CRELD2, PDIA6, DPAGT1, CDK2AP2, NUCB2, and SEC61A1, eight of which localized to the upregulated Module 2. Downregulated hub genes included LUM, FBN1, SPARC, LAMA4, EFEMP2, NID2, AEBP1, PDGFRB, CTSK, and COL4A2, several of which clustered within downregulated Modules 2 and 3 (Figure 1E; Supplementary Figure S1). In fusion-negative tumors, PPI networks constructed from 843 upregulated and 455 downregulated fusion-negative specific DEGs also resolved into three significant modules. MCC analysis highlighted upregulated hubs PBK, MELK, KIF23, RAD54L, CDC25C, TTK, SPC25, CDKN3, TRIP13, and NCAPG2, as well as downregulated hubs MORN5, DNAI2, DNAAF6, TEKT2, LRRC71, DTHD1,TEKT1, ZBBX, MAP3K19, and DNAI1, all of which were associated with Module 1. Expression patterns of these hub genes are summarized in Table 1.

**Table 1.**
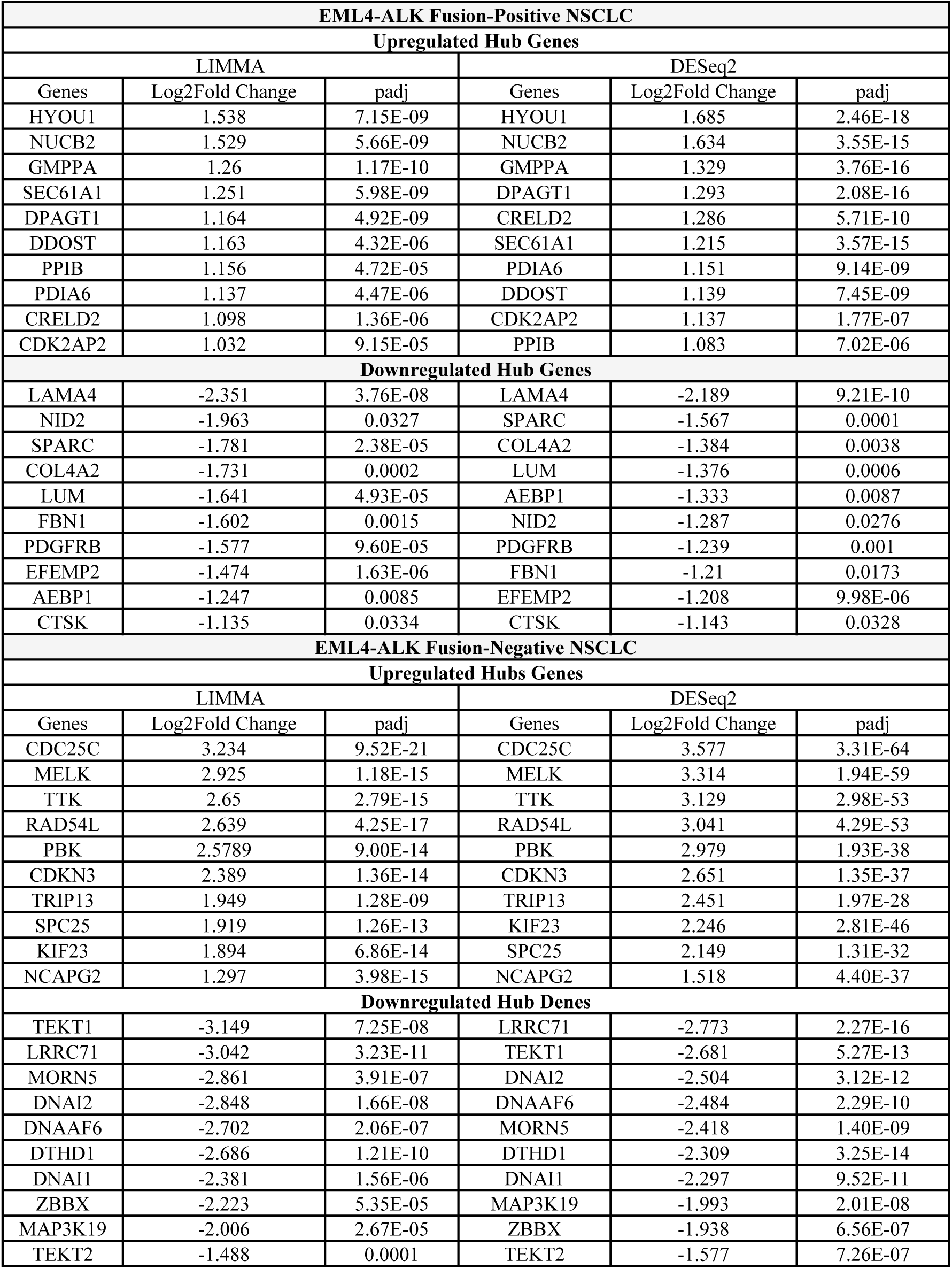
List of the identified hub DEGs in fusion-positive and fusion-negative NSCLC using LIMMA and DESeq2 methods.

| EML4-ALK Fusion-Positive NSCLC |  |  |  |  |  |
| --- | --- | --- | --- | --- | --- |
| Upregulated Hub Genes |  |  |  |  |  |
| LIMMA |  |  | DESeq2 |  |  |
| Genes | Log2Fold Change | padj | Genes | Log2Fold Change | padj |
| HYOU1 | 1.538 | 7.15E-09 | HYOU1 | 1.685 | 2.46E-18 |
| NUCB2 | 1.529 | 5.66E-09 | NUCB2 | 1.634 | 3.55E-15 |
| GMPPA | 1.26 | 1.17E-10 | GMPPA | 1.329 | 3.76E-16 |
| SEC61A1 | 1.251 | 5.98E-09 | DPAGT1 | 1.293 | 2.08E-16 |
| DPAGT1 | 1.164 | 4.92E-09 | CRELD2 | 1.286 | 5.71E-10 |
| DDOST | 1.163 | 4.32E-06 | SEC61A1 | 1.215 | 3.57E-15 |
| PPIB | 1.156 | 4.72E-05 | PDIA6 | 1.151 | 9.14E-09 |
| PDIA6 | 1.137 | 4.47E-06 | DDOST | 1.139 | 7.45E-09 |
| CRELD2 | 1.098 | 1.36E-06 | CDK2AP2 | 1.137 | 1.77E-07 |
| CDK2AP2 | 1.032 | 9.15E-05 | PPIB | 1.083 | 7.02E-06 |
| Downregulated Hub Genes |  |  |  |  |  |
| LAMA4 | -2.351 | 3.76E-08 | LAMA4 | -2.189 | 9.21E-10 |
| NID2 | -1.963 | 0.0327 | SPARC | -1.567 | 0.0001 |
| SPARC | -1.781 | 2.38E-05 | COL4A2 | -1.384 | 0.0038 |
| COL4A2 | -1.731 | 0.0002 | LUM | -1.376 | 0.0006 |
| LUM | -1.641 | 4.93E-05 | AEBP1 | -1.333 | 0.0087 |
| FBN1 | -1.602 | 0.0015 | NID2 | -1.287 | 0.0276 |
| PDGFRB | -1.577 | 9.60E-05 | PDGFRB | -1.239 | 0.001 |
| EFEMP2 | -1.474 | 1.63E-06 | FBN1 | -1.21 | 0.0173 |
| AEBP1 | -1.247 | 0.0085 | EFEMP2 | -1.208 | 9.98E-06 |
| CTSK | -1.135 | 0.0334 | CTSK | -1.143 | 0.0328 |
| EML4-ALK Fusion-Negative NSCLC |  |  |  |  |  |
| Upregulated Hubs Genes |  |  |  |  |  |
| LIMMA |  |  | DESeq2 |  |  |
| Genes | Log2Fold Change | padj | Genes | Log2Fold Change | padj |
| CDC25C | 3.234 | 9.52E-21 | CDC25C | 3.577 | 3.31E-64 |
| MELK | 2.925 | 1.18E-15 | MELK | 3.314 | 1.94E-59 |
| TTK | 2.65 | 2.79E-15 | TTK | 3.129 | 2.98E-53 |
| RAD54L | 2.639 | 4.25E-17 | RAD54L | 3.041 | 4.29E-53 |
| PBK | 2.5789 | 9.00E-14 | PBK | 2.979 | 1.93E-38 |
| CDKN3 | 2.389 | 1.36E-14 | CDKN3 | 2.651 | 1.35E-37 |
| TRIP13 | 1.949 | 1.28E-09 | TRIP13 | 2.451 | 1.97E-28 |
| SPC25 | 1.919 | 1.26E-13 | KIF23 | 2.246 | 2.81E-46 |
| KIF23 | 1.894 | 6.86E-14 | SPC25 | 2.149 | 1.31E-32 |
| NCAPG2 | 1.297 | 3.98E-15 | NCAPG2 | 1.518 | 4.40E-37 |
| Downregulated Hub Denes |  |  |  |  |  |
| TEKT1 | -3.149 | 7.25E-08 | LRRC71 | -2.773 | 2.27E-16 |
| LRRC71 | -3.042 | 3.23E-11 | TEKT1 | -2.681 | 5.27E-13 |
| MORN5 | -2.861 | 3.91E-07 | DNAI2 | -2.504 | 3.12E-12 |
| DNAI2 | -2.848 | 1.66E-08 | DNAAF6 | -2.484 | 2.29E-10 |
| DNAAF6 | -2.702 | 2.06E-07 | MORN5 | -2.418 | 1.40E-09 |
| DTHD1 | -2.686 | 1.21E-10 | DTHD1 | -2.309 | 3.25E-14 |
| DNAI1 | -2.381 | 1.56E-06 | DNAI1 | -2.297 | 9.52E-11 |
| ZBBX | -2.223 | 5.35E-05 | MAP3K19 | -1.993 | 2.01E-08 |
| MAP3K19 | -2.006 | 2.67E-05 | ZBBX | -1.938 | 6.56E-07 |
| TEKT2 | -1.488 | 0.0001 | TEKT2 | -1.577 | 7.26E-07 |

### Fusion-positive NSCLC is marked by activation of ER proteostasis and glycolytic programs with suppression of adhesion and immune signaling

Gene Ontology (GO) analysis of upregulated DEGs implicated intracellular signal transduction, cytokine-mediated signaling, cell proliferation, regulation of TOR signaling, apoptotic processes, and protein folding and binding (Figure 1F; Supplementary Table S1). KEGG pathway analysis revealed enrichment of glycolysis/gluconeogenesis, protein processing in the endoplasmic reticulum, N-glycan biosynthesis, and p53 signaling (Figure 1G; Supplementary Table S2). Notably, several hub genes mapped directly to these pathways, including DDOST (ER processing and N-glycan biosynthesis), DPAGT1 (N-glycan biosynthesis), and PDIA6 (protein folding). In contrast, downregulated genes were enriched for cell–cell adhesion, actin cytoskeleton regulation, protein localization, immune responses, and cell-surface receptor signaling (Figure 1F; Supplementary Table S1), with KEGG under-representation of T-cell receptor signaling, cytokine–cytokine receptor interaction, calcium signaling, general cancer pathways, and chemical carcinogenesis–receptor activation (Figure 1G; Supplementary Table S2).

### Fusion-negative NSCLC shows enrichment of cell-cycle and DNA-repair programs with repression of ciliary and microtubule-associated pathways

Upregulated DEGs in fusion-negative tumors were enriched for cell division, chromosome segregation, regulation of gene expression, heterochromatin assembly, ATPase activity, and DNA repair (Figure 1F; Supplementary Table S1). KEGG analysis highlighted enrichment of cell-cycle regulation, homologous recombination, microRNAs in cancer, ATP-dependent chromatin remodeling, and transcriptional misregulation in cancer (Figure 1G; Supplementary Table S2). Hub genes TTK, TRIP13, and CDC25C mapped to cell-cycle pathways, RAD54L to homologous recombination, and KIF23 to motor-protein activity, with KIF23 and CDC25C also participating in microRNA-associated cancer pathways. Conversely, downregulated DEGs were enriched for cilium movement and assembly, microtubule-based transport, negative regulation of Wnt signaling, calcium-mediated signaling, and positive regulation of gene expression (Figure 1F; Supplementary Table S1). Corresponding KEGG pathway depletion included motor-protein pathways, neurodegenerative modules, drug metabolism via cytochrome P450, tyrosine metabolism, and lipid and atherosclerosis pathways (Figure 1G; Supplementary Table S2). The ciliary hub genes DNAI1 and DNAI2 contributed prominently to these suppressed functional categories.

### Pathway-level perturbations stratified by fusion status

Gene set enrichment analysis (GSEA) of ranked expression profiles comparing fusion-positive and fusion-negative tumors revealed positive enrichment of amino- and nucleotide-sugar metabolism, together with protein export in fusion-positive NSCLC, including β-alanine,phenylalanine, propanoate, pyruvate, fatty-acid, histidine, arginine–proline, and butanoate metabolism, together with protein export (Figure 2A). Leading-edge genes such as GMPPA, involved in amino- and nucleotide-sugar metabolism, and SEC61A1, a core component of the ER translocon and protein-export machinery, anchored these signals, linking transcriptional upregulation to glycan biosynthesis and secretory pathways (Figure 2B). In contrast, pathways downregulated in fusion-positive relative to fusion-negative tumors included cytokine–cytokine receptor interaction, ECM–receptor interaction, calcium signaling, focal adhesion, and Toll-like receptor signaling (Figure 2C). Projection of hub genes onto these pathways implicated PDGFRB (cytokine receptor, calcium, gap junction, and cancer signaling), ECM components LAMA4 and COL4A2 (ECM–receptor interaction, focal adhesion, cancer pathways), and CTSK (Toll-like receptor signaling).

**Figure 2.**
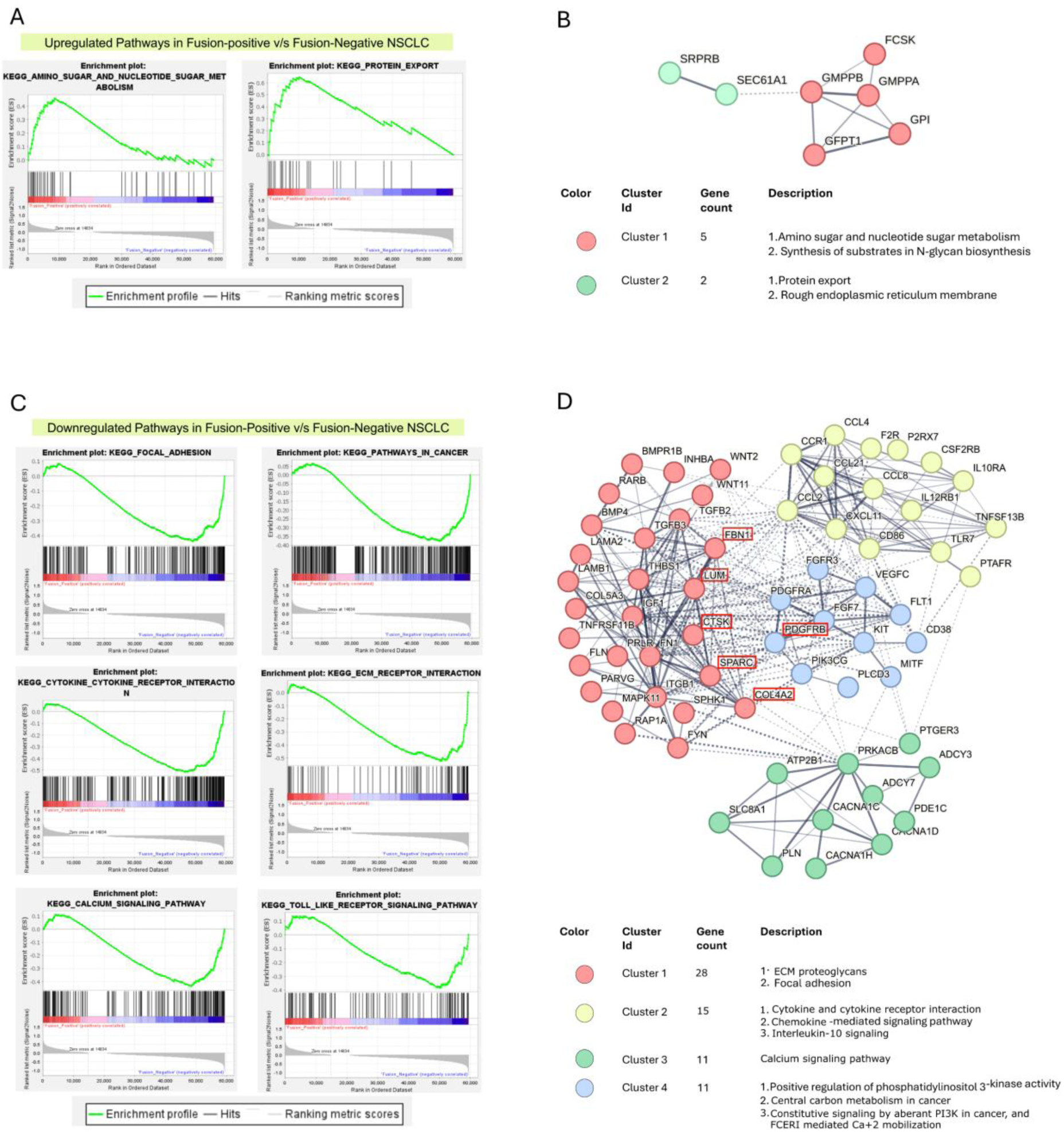
GSEA uncovers metabolic induction and immune-stromal attenuation in EML4-ALK fusion-positive NSCLC. (A) Positively enriched pathways in fusion-positive versus fusion-negative tumors. (B) PPI network of fusion-positive exclusive up-regulated genes annotated by the pathways. (C) Negatively enriched pathways in fusion-positive versus fusion-negative tumors, including cytokine-receptor, ECM-receptor, calcium signaling, gap junction, focal adhesion, cancer pathways, and Toll-like receptor signaling. (D) PPI network of fusion-positive exclusive down-regulated genes. MicroRNA hsa-miR-3065, predicted to target six hub genes (PDGFRB, CTSK, COL4A2, SPARC, FBN1, LUM) shown in red boxes. Dotted contours denote k-means clusters capturing pathway interconnections.

### Network organization of GSEA-derived signals

To further resolve pathway organization, k-means clustering of pathway-annotated PPI networks partitioned GSEA-enriched signals into discrete functional modules. In the upregulated network (Figure 2B), Cluster 1, comprising FCSK, GMPPB, GMPPA, GPI, and GFPT1, captured amino- and nucleotide-sugar metabolism and N-glycan substrate synthesis, whereas Cluster 2, including SEC61A1 and SRPRB, represented ER-associated protein export. The inter-cluster connectivity indicated coordinated coupling between hexosamine/glycan flux and ER translocation.

In the downregulated network (Figure 2D), Cluster 1 corresponded to focal adhesion and ECM proteoglycan pathways; Cluster 2 to cytokine–cytokine receptor interaction, chemokine-mediated signaling, and IL-10 signaling; Cluster 3 to calcium signaling; and Cluster 4 to positive regulation of PI3K signaling, aberrant PI3K signaling in cancer, and VEGF-associated interactions involving VEGFC, FLT1, FGFR3, KIT, PDGFRA, and PIK3CG. Inter-cluster connectivity between adhesion-related pathways in Cluster 1 and PI3K/VEGF signaling in Cluster 4 indicates an ECM–PI3K crosstalk. Notably, five downregulated hub genes—CTSK, COL4A2, SPARC, FBN1, and LUM—mapped to Cluster 1, whereas PDGFRB mapped to Cluster 4. These six hub genes were all predicted targets of the upregulated microRNA hsa-miR-3065.

### Comparative miRNA expression profiling and target regulation stratified by fusion status

We analyzed miRNA expression profiles in EML4–ALK fusion-positive and fusion-negative NSCLC relative to normal lung tissue. Differentially expressed miRNAs (DEMs) were identified using DESeq2 and Limma and visualized using volcano plots (Figure 3B; Table 2). Intersecting results from both methods identified eight upregulated and eight downregulated DEMs in fusion-positive tumors, and 83 upregulated and 65 downregulated DEMs in fusion-negative tumors (Figure 3C). MicroRNA hsa-miR-3065 was the only upregulated DEM unique to fusion-positive cases, with no fusion-positive exclusive downregulated miRNAs detected, whereas fusion-negative tumors harbored 76 exclusively upregulated and 57 exclusively downregulated DEMs (Figure 3C; Supplementary Table S3). To associate miRNAs with putative mRNA targets, we queried mirDIP using medium-confidence criteria (consensus across at least three databases). In fusion-positive tumors, hsa-miR-3065-3p/5p was predicted to target 398 downregulated genes, including six downregulated hub genes identified in the PPI analysis: PDGFRB, CTSK, COL4A2, SPARC, FBN1, and LUM (Figure 3D).

**Figure 3.**
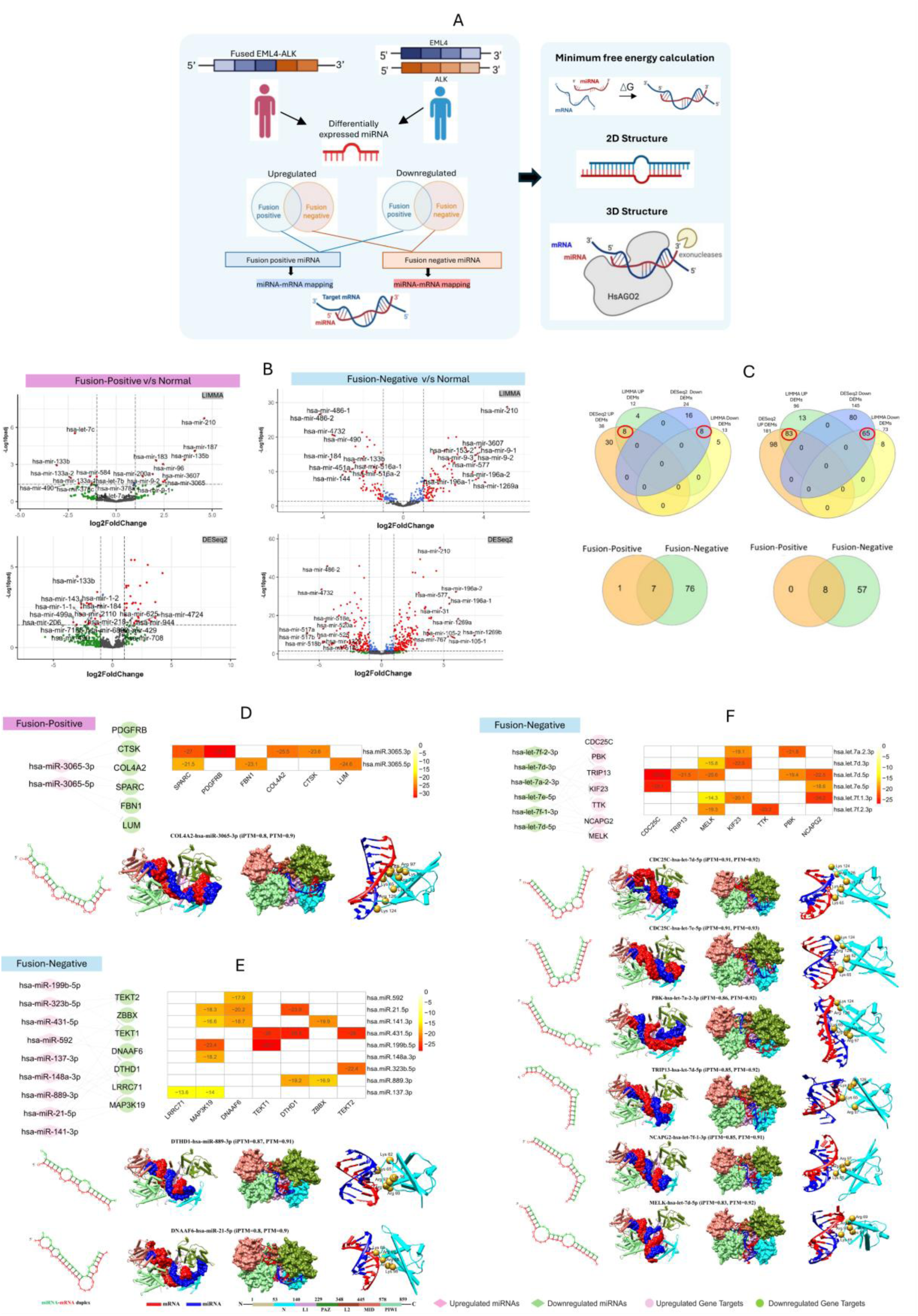
miRNA expression and target regulation stratified by EML4-ALK fusion status. (A) Schematic of the analysis pipeline: DEM discovery (DESeq2, limma), intersected sets, MFE screening, and 2D/3D modeling of AGO2-bound miRNA-mRNA complexes. (B) Volcano plots of DEMs for fusion-positive vs normal and fusion-negative vs normal. (C) Venn diagrams: method intersections for up- and down-regulated DEMs and subtype-exclusive DEMs (fusion-positive vs fusion-negative). (D) Fusion-positive network: hsa-miR-3065-3p/5p targeting of down-regulated hub genes (PDGFRB, CTSK, COL4A2, SPARC, FBN1, LUM), with MFE heatmap and representative 2D/3D complexes (AGO2 compatible). (E) Fusion-negative network: top up-regulated miRNAs predicted to target down-regulated hubs (TEKT2, TEKT1, ZBBX, DNAAF6, DTHD1, LRRC71, MAP3K19), with MFE heatmap and structures. (F) Fusion-negative network: down-regulated let-7 miRNAs targeting up-regulated hubs (CDC25C, PBK, TRIP13, KIF23, TTK, NCAPG2, MELK), with MFE heatmap and 2D/3D models. Heatmaps show RNAhybrid minimum free energy (kcal/mol); 3D models were selected at MFE ≤ −18 kcal mol⁻¹ and ipTM ≥ 0.8 / pTM ≥ 0.9.

**Table 2.** List of the identified key DEMs in fusion-positive and fusion-negative NSCLC.

| EML4-ALK Fusion-Positive NSCLC |  |  |  |  |  |
| --- | --- | --- | --- | --- | --- |
| Upregulated MicroRNA |  |  |  |  |  |
| LIMMA |  |  | DESeq2 |  |  |
| miRNA | Log2Fold | padj | miRNA | Log2Fold | padj |
| hsa-mir-3065 | 2.443 | 0.0251 | hsa-mir-3065 | 2.991 | 7.39E-06 |
| EML4-ALK Fusion-Negative NSCLC |  |  |  |  |  |
| Upregulated MicroRNA |  |  |  |  |  |
| LIMMA |  |  | DESeq2 |  |  |
| miRNA | Log2Fold | padj | miRNA | Log2Fold | padj |
| hsa-mir-323b | 2.584 | 8.42E-07 | hsa-mir-323b | 3.818 | 3.82E-11 |
| hsa-mir-21 | 2.425 | 3.85E-10 | hsa-mir-137 | 3.694 | 2.14E-10 |
| hsa-mir-141 | 1.807 | 1.27E-08 | hsa-mir-889 | 2.496 | 1.31E-12 |
| hsa-mir-148a | 1.712 | 6.14E-10 | hsa-mir-431 | 2.427 | 9.02E-09 |
| hsa-mir-592 | 1.5109 | 7.04E-06 | hsa-mir-21 | 1.65 | 1.53E-13 |
| hsa-mir-199b | 1.426 | 3.17E-08 | hsa-mir-141 | 1.627 | 1.09E-14 |
| hsa-mir-137 | 1.317 | 0.0001 | hsa-mir-148a | 1.593 | 7.05E-20 |
| hsa-mir-431 | 1.247 | 0.0074 | hsa-mir-592 | 1.568 | 1.13E-06 |
| hsa-mir-889 | 1.148 | 0.0043 | hsa-mir-199b | 1.43 | 8.31E-17 |
| Downregulated MicroRNA |  |  |  |  |  |
| hsa-let-7a-2 | -1.821 | 8.39E-21 | hsa-let-7d | -2.388 | 5.54E-22 |
| hsa-let-7f-2 | -1.619 | 3.57E-05 | hsa-let-7f-2 | -2.038 | 1.41E-17 |
| hsa-let-7f-1 | -1.614 | 4.04E-05 | hsa-let-7f-1 | -2.038 | 1.70E-17 |
| hsa-let-7e | -1.44 | 4.26E-12 | hsa-let-7a-2 | -1.783 | 1.46E-46 |
| hsa-let-7d | -1.177 | 1.71E-10 | hsa-let-7e | -1.339 | 6.48E-17 |

In fusion-negative tumors, 76 upregulated DEMs were predicted to target 440 downregulated genes, whereas 57 downregulated DEMs mapped to 463 upregulated genes. The upregulated miRNAs (hsa-miR-137-3p, hsa-miR-148a-3p, hsa-miR-199b-5p, hsa-miR-141-3p, hsa-miR-21-5p, hsa-miR-592, hsa-miR-431-5p, hsa-miR-889-3p, and hsa-miR-323b-5p) collectively targeted downregulated hub genes TEKT2, TEKT1, ZBBX, DNAAF6, DTHD1, LRRC71, and MAP3K19 (Figure 3E). Conversely, downregulated members of the let-7 family (let-7f-2-3p, let-7d-3p, let-7a-2-3p, let-7e-5p, let-7f-1-3p, let-7d-5p) were predicted to target upregulated hub genes CDC25C, PBK, TRIP13, KIF23, TTK, NCAPG2, and MELK (Figure 3F).

### Structure-guided analysis of miRNA–mRNA recognition

Minimum free-energy (MFE) screening (≤ −18 kcal mol⁻¹) was applied to prioritize miRNA–mRNA pairs for structural modeling. In fusion-positive lung adenocarcinoma, four candidate miRNA–mRNA pairs, selected from PDGFRB, COL4A2, FBN1, and LUM, were subjected to RNAhybrid-based secondary-structure prediction and AlphaFold-based three-dimensional modeling. Two of these pairs, COL4A2, and FBN1 achieved ipTM ≥ 0.8 and pTM ≥ 0.9 (Table 3), and their predicted duplex structures are shown in Supplementary Figures S3–S5. Notably, the COL4A2–hsa-miR-3065-3p duplex adopted an AGO2-compatible conformation, occupying the central channel and contacting the N, L1, PAZ, L2, MID, and PIWI domains ^58^. Interface analysis within a 6Å threshold highlighted interactions with conserved basic residues (Lys62, Lys65, Arg68, Arg69, Arg72, Arg97, Lys98, Lys124, Arg126), consistent with AGO2-mediated positioning of the miR-3065 seed on the COL4A2 3′-UTR (Figure 3D).

**Table 3.** Interface predicted template modelling (iPTM) and predicted template modeling (PTM) scores for 3D structures of influential miRNA-mRNA pairs in fusion positive NSCLC cases.

| Fusion-Positive NSCLC |  |  |  |
| --- | --- | --- | --- |
| mRNA | miRNA | iPTM score | PTM score |
| COL4A2 | hsa-miR-3065-3p | 0.8 | 0.9 |
| FBN1 | hsa-miR-3065-5p | 0.8 | 0.9 |
| PDGFRB | hsa-miR-3065-3p | 0.72 | 0.88 |
| LUM | hsa-miR-3065-5p | 0.69 | 0.91 |
| Fusion-Negative NSCLC |  |  |  |
| mRNA | miRNA | iPTM score | PTM score |
| CDC25C | hsa-let-7d-5p | 0.91 | 0.92 |
| CDC25C | hsa-let-7e-5p | 0.91 | 0.93 |
| DTHD1 | hsa-miR-889-3p | 0.87 | 0.91 |
| PBK | hsa-let-7a-2-3p | 0.86 | 0.92 |
| TRIP13 | hsa-let-7d-5p | 0.85 | 0.92 |
| NCAPG2 | hsa-let-7f-1-3p | 0.85 | 0.91 |
| MELK | hsa-let-7d-5p | 0.83 | 0.92 |
| DNAAF6 | hsa-miR-21-5p | 0.8 | 0.9 |
| KIF23 | hsa-let-7d-3p | 0.8 | 0.87 |
| PBK | hsa-let-7d-5p | 0.8 | 0.89 |
| DNAAF6 | hsa-miR-592 | 0.79 | 0.92 |
| ZBBX | hsa-miR-889-3p | 0.78 | 0.88 |
| TTK | hsa-let-7f-2-3p | 0.78 | 0.9 |
| NCAPG2 | hsa-let-7d-5p | 0.77 | 0.92 |
| NCAPG2 | hsa-let-7e-5p | 0.77 | 0.9 |
| DNAAF6 | hsa-miR-141-3p | 0.64 | 0.9 |
| ZBBX | hsa-miR-141-3p | 0.6 | 0.89 |

In fusion-negative tumors, 27 candidate miRNA–mRNA pairs were screened, of which 10 satisfied the ipTM ≥ 0.8 and pTM ≥ 0.9 criteria. Among these, CDC25C–let-7d-5p, CDC25C–let-7e-5p, DTHD1–miR-889-3p, PBK–let-7a-2-3p, TRIP13–let-7d-5p, NCAPG2–let-7f-1-3p, MELK–let-7d-5p, and DNAAF6–miR-21-5p formed AGO2-compatible complexes, with conserved basic residues in the AGO2 N domain stabilizing seed-region positioning on the target 3′-UTRs (Figure 3E–F).

### EML4/ALK-targeting miRNAs and a fusion-driven sponge mechanism

To investigate post-transcriptional regulation linked directly to the EML4–ALK fusion, we first cataloged miRNAs predicted to bind the EML4 and ALK parental transcripts. In silico screening identified 304 EML4-targeting and 40 ALK-targeting miRNAs, of which TCGA lung miRNA-seq data indicated that 98 and 12, respectively, were expressed in lung tissue. These expressed miRNAs were then mapped onto fusion and non-fusion-specific DEG sets. Collectively, the 98 EML4-targeting miRNAs were predicted to regulate 623 downregulated genes, whereas the 12 ALK-targeting miRNAs were associated with 179 upregulated genes in fusion-positive NSCLC, yielding 5,554 EML4–miRNA–gene regulatory pairs (Supplementary Figure S2).

Network projection revealed convergence of EML4-targeting hub miRNAs on six downregulated hub genes previously identified in the PPI analysis—FBN1, LAMA4, NID2, COL4A2, PDGFRB, and SPARC whereas among ALK-targeting hub genes, only HYOU1 emerged as a predicted target, mediated by hsa-miR-6501-3p (Figure 4B). Minimum free-energy estimates derived from RNAhybrid supported high-affinity interactions (≤ −18 kcal mol⁻¹), including those involving members of the miR-29 family (hsa-miR-29c-3p, hsa-miR-29a-3p), hsa-miR-152-3p, hsa-miR-199b-3p, hsa-miR-627-5p, and hsa-miR-6501-3p with both hub genes and the parental transcripts (Figure 4C).

**Figure 4.**
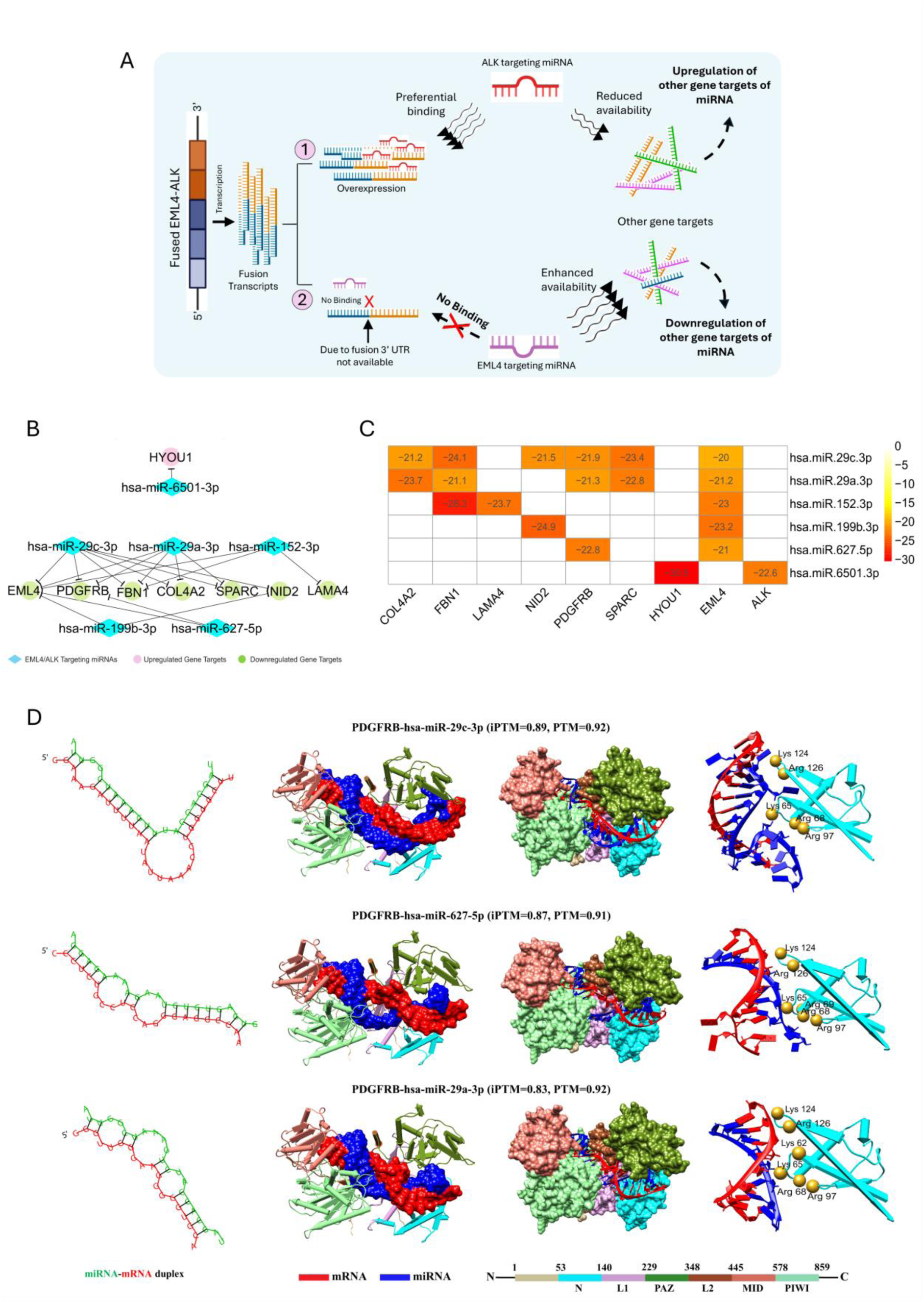
Fusion-driven sponging of EML4/ALK-targeting miRNAs and downstream regulatory consequences. (A) Schematic model of fusion-driven miRNA sponging and downstream regulation. Cartoon depicting how the EML4-ALK fusion can rewire post-transcriptional control: (1) overexpressed fusion transcripts preferentially engage ALK-targeting miRNAs, reducing their availability for other mRNA targets and favoring up-regulation of those genes; (2) loss/inaccessibility of the native EML4 3′UTR limits binding of EML4-targeting miRNAs, increasing the free miRNA pool and thereby enhancing repression of their alternative targets, yielding down-regulation. (B) miRNA-mRNA network highlighting EML4/ALK-targeting hub miRNAs (diamonds) and their gene targets (pink, up-regulated; green, down-regulated). Notable edges include hsa-miR-6501-3p to HYOU1 and EML4-targeting miRNAs (miR-29c-3p, miR-29a-3p, miR-152-3p, miR-199b-3p, miR-627-5p) to PDGFRB, FBN1, COL4A2, SPARC, NID2, LAMA4. (C) Heat map of RNAhybrid minimum free energies (kcal mol⁻¹) for selected miRNA-mRNA pairs, including EML4 and ALK. (D) Representative duplexes-PDGFRB-miR-29c-3p, PDGFRB-miR-627-5p, PDGFRB-miR-29a-3p-with 2D structures (left) and AlphaFold-based 3D models bound to AGO2 (center/right). Reported ipTM/pTM values indicate model confidence; residue labels mark conserved basic contacts at the duplex-AGO2 interface.

To evaluate structural plausibility, representative miRNA–mRNA duplexes were modeled in three dimensions. Three PDGFRB–miRNA complexes—PDGFRB–hsa-miR-29c-3p (ipTM = 0.89, pTM = 0.92), PDGFRB–hsa-miR-627-5p (ipTM = 0.87, pTM = 0.91), and PDGFRB–hsa-miR-29a-3p (ipTM = 0.83, pTM = 0.92)—adopted AGO2-compatible conformations, with duplexes spanning the N, L1, PAZ, L2, MID, and PIWI domains and engaging conserved basic residues such as Lys62, Arg68, Arg97, Lys124, and Arg126 (Figure 4D; Table 4).

**Table 4.** Prediction scores for 3D structures of EML4 and ALK targeting miRNA pairs.

| mRNA | EML4 and ALK targeting miRNA | iPTM score* | PTM score# |
| --- | --- | --- | --- |
| PDGFRB | hsa-miR-29c-3p | 0.89 | 0.92 |
| COL4A2 | hsa-miR-29a-3p | 0.89 | 0.93 |
| PDGFRB | hsa-miR-627-5p | 0.87 | 0.91 |
| PDGFRB | hsa-miR-29a-3p | 0.83 | 0.92 |
| NID2 | hsa-miR-29c-3p | 0.79 | 0.89 |
| EML4 | hsa-miR-627-5p | 0.79 | 0.9 |
| LAMA4 | hsa-miR-152-3p | 0.78 | 0.92 |
| ALK | hsa-miR-6501-3p | 0.77 | 0.9 |
| HYOU1 | hsa-miR-6501-3p | 0.7 | 0.87 |
| NID2 | hsa-miR-199b-3p | 0.69 | 0.9 |
| EML4 | hsa-miR-29a-3p | 0.61 | 0.88 |
| FBN1 | hsa-miR-152-3p | 0.59 | 0.89 |
| EML4 | hsa-miR-29c-3p | 0.56 | 0.88 |
| COL4A2 | hsa-miR-29c-3p | 0.53 | 0.89 |
| EML4 | hsa-miR-152-3p | 0.53 | 0.89 |
| FBN1 | hsa-miR-29c-3p | 0.5 | 0.89 |
| EML4 | hsa-miR-199b-3p | 0.5 | 0.89 |
| FBN1 | hsa-miR-29a-3p | 0.31 | 0.87 |

\*iPTM =interface predicted template modelling; #PTM =predicted template modeling

Together with the schematic in Figure 4A, these findings support a fusion-driven sponge model in which preferential engagement of ALK-targeting miRNAs by the overexpressed fusion transcript reduces their availability for alternative targets, contributing to upregulation of those genes, while loss or masking of the native EML4 3′-UTR increases the effective pool of EML4-targeting miRNAs, enhancing repression of their downstream targets. This regulatory architecture provides a model depicting mechanistic link between the fusion event and coordinated repression of ECM and adhesion hubs alongside selective modulation of stress-response genes in fusion-positive NSCLC.

### Quantitative assessment of stromal and immune cell infiltration in fusion-defined NSCLC tumor microenvironments

To characterize the tumor immune microenvironment (TIME) in EML4–ALK fusion-positive and fusion-negative NSCLC, we quantitatively evaluated stromal, immune, and ESTIMATE scores for each tumor sample. While immune and ESTIMATE scores did not differ significantly between the two groups, stromal scores were significantly lower in fusion-positive NSCLC compared with fusion-negative tumors (Figure 5A). To further delineate the cellular composition of the TIME, we applied the MCP-counter immune deconvolution to estimate infiltration levels of key stromal and immune cell populations, including T cells, CD8⁺ T cells, NK cells, B cells, macrophages, myeloid dendritic cells, neutrophils, endothelial cells, cytotoxicity scores, and cancer-associated fibroblasts (CAFs). Among these populations, CAFs were significantly less abundant in fusion-positive NSCLC samples relative to fusion-negative tumors (Figure 5B), indicating distinct stromal architectures associated with fusion status.

**Figure 5.**
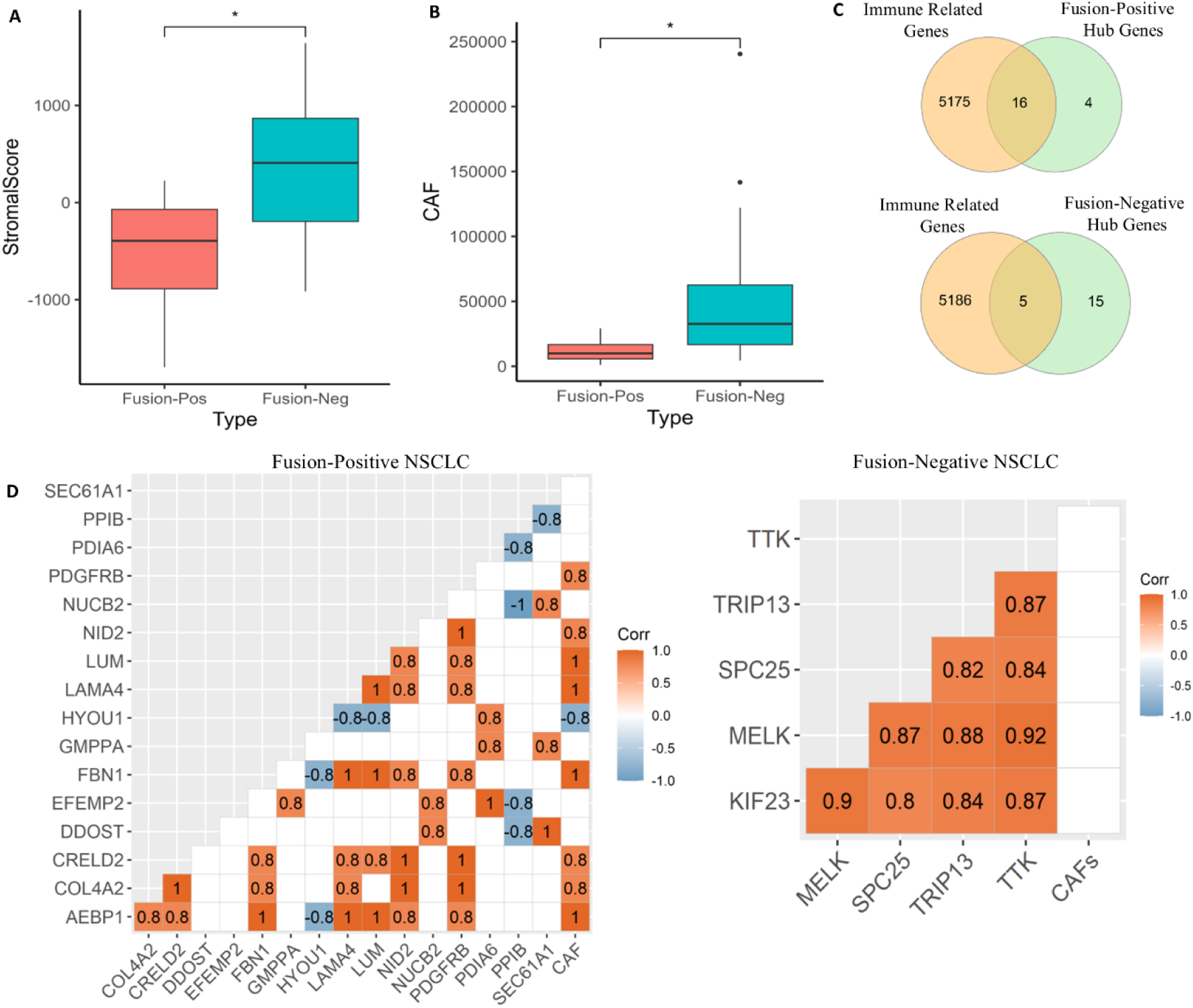
Stromal depletion and CAF signatures differ by EML4-ALK fusion status. (A) Boxplots of StromalScore (ESTIMATE) for fusion-positive and fusion-negative tumors; boxes show median and interquartile range, whiskers 1.5× IQR; asterisks denote P < 0.05. (B) MCP-counter-derived CAF abundance, displayed as in (A). (C) Venn overlaps between immune-related gene set and subtype hub genes: intersection counts highlight 16 immune-related hubs in fusion-positive tumors and 5 in fusion-negative tumors. (D) Spearman correlation heat maps. Left: fusion-positive NSCLC showing associations among CAFs and ECM/adhesion hubs alongside upregulated hubs. Right: fusion-negative NSCLC showing strong inter-correlation among proliferative hubs with minimal correlation to CAFs.

To explore molecular factors underlying differential CAF infiltration, we examined a curated set of 5,191 immune-associated genes and identified differentially expressed hub immune-related genes in both cohorts (Figure 5C; Supplementary Table S4). We then performed Spearman correlation analyses between CAF infiltration levels and gene expression profiles to assess gene–CAF associations (Figure 5D). In fusion-positive NSCLC, strong positive correlations were observed between CAF infiltration and several downregulated hub genes, including PDGFRB, NID2, LUM, LAMA4, FBN1, COL4A2, and AEBP1. These genes also exhibited high inter-correlation, consistent with a coordinated regulatory program. An upregulated hub gene HYOU1 displayed a strong negative correlation with CAF levels as well as with the CAF-associated hub genes including LAMA4, LUM, AEBP1 and FBN1.

In contrast, in fusion-negative NSCLC, the upregulated hub genes SPC25, TRIP13, MELK, KIF23, and TTK showed no significant association with CAF infiltration but demonstrated strong inter-correlation within the TIME. These patterns suggest a distinct regulatory architecture in fusion-negative tumors, in which oncogenic signaling may proceed largely independently of CAF-mediated stromal interactions. Overall, these analyses indicate that the CAF landscape differs substantially between EML4–ALK fusion-positive and fusion-negative NSCLC, with subtype-specific gene expression signatures underlying this divergence.

### ROC curve analysis identifies potential biomarkers for fusion-positive and fusion-negative NSCLC

To assess the diagnostic potential of selected differentially expressed genes (DEGs) and miRNAs, we performed receiver operating characteristic (ROC) curve analyses on hub genes and miRNAs associated with extracellular matrix organization, cell adhesion, and CAF-related processes. ROC curves were generated for nine hub genes identified in fusion-positive NSCLC: SPARC, PDGFRB, FBN1, COL4A2, CTSK, LUM, LAMA4, NID2, and HYOU1. Six of these genes exceeded the diagnostic threshold (AUC > 0.7), with LAMA4 showing the highest predictive value (AUC = 0.97), followed by HYOU1 (AUC = 0.95), LUM (AUC = 0.83), SPARC (AUC = 0.83), PDGFRB (AUC = 0.73), and COL4A2 (AUC = 0.71) (Figure 6A). We also evaluated hsa-miR-3065, the most prominently upregulated miRNA in fusion-positive NSCLC, which demonstrated strong discriminatory performance with an AUC of 0.91 (Figure 6B). Notably, hsa-miR-3065 and four of its downregulated target genes including PDGFRB, COL4A2, SPARC, and LUM exhibited consistent predictive accuracy, supporting their potential utility as biomarkers for fusion-positive NSCLC. Similarly, ROC analysis was performed for 14 key hub genes in fusion-negative NSCLC: LRRC71, MAP3K19, DNAAF6, TEKT1, DTHD1, ZBBX, TEKT2, CDC25C, TRIP13, MELK, KIF23, TTK, PBK, and NCAPG2. Eleven of these genes exceeded the AUC threshold of 0.7 (Figure 6C), whereas MAP3K19, ZBBX, and TEKT2 did not meet the predictive criterion.

**Figure 6.**
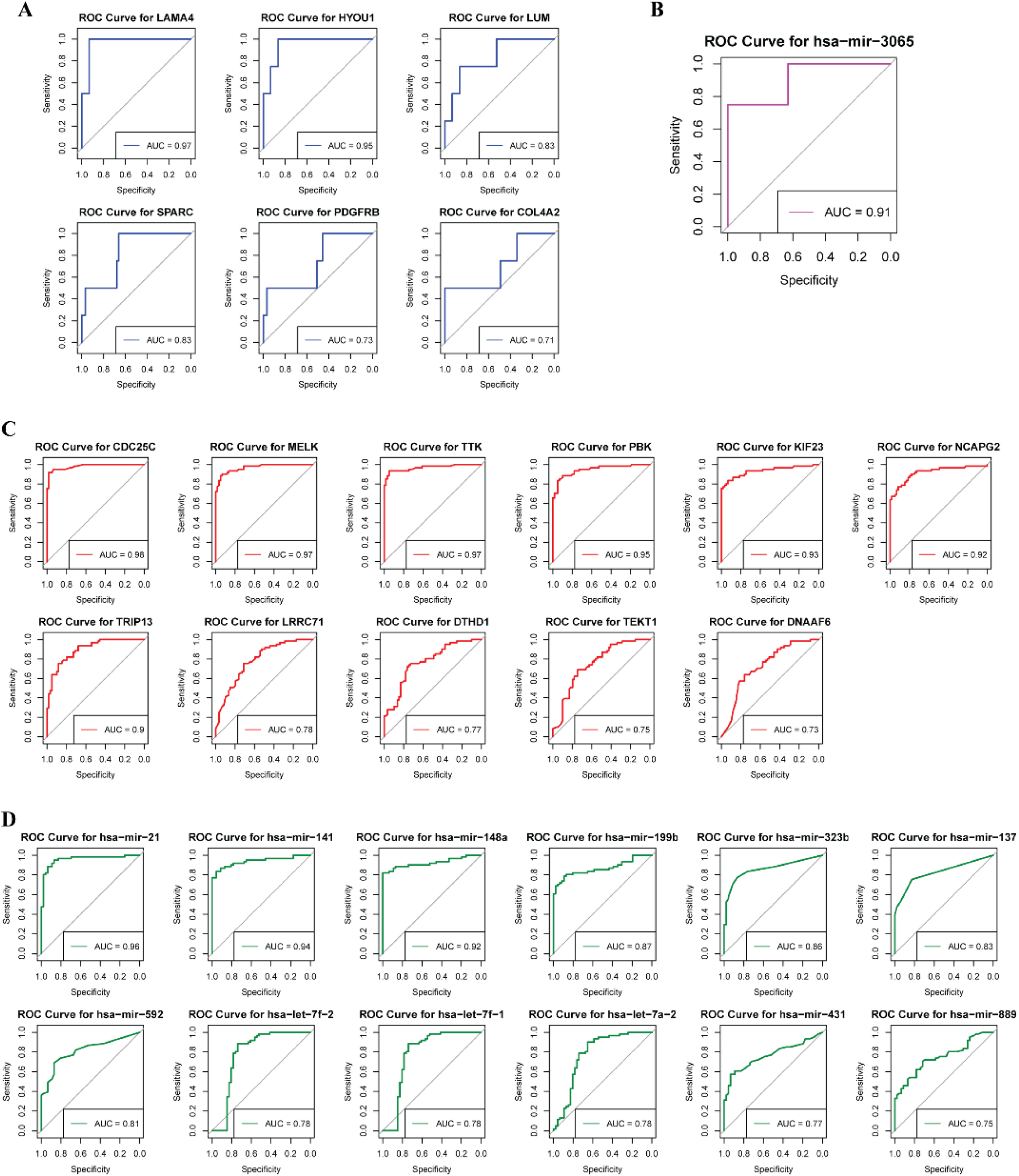
The ROC curves of identified hub genes and miRNAs. The analysis evaluates the predictive value of the biomarker genes and miRNAs in NSCLC. The figure depicts AUC value and ROC of (A & B) biomarker genes and miRNAs of fusion-positive NSCLC cases and (C & D) biomarker genes and miRNA in fusion-negative NSCLC cases. The cut-off value of AUC ≥0.7 was chosen for a good predictor.

Analysis of fusion-negative–specific miRNAs showed that 12 of 14 miRNAs achieved AUC values above 0.7. Among these, hsa-miR-21 emerged as the strongest predictor (AUC = 0.96), followed by hsa-miR-141 (AUC = 0.94), hsa-miR-148a (AUC = 0.92), and hsa-miR-199b (AUC = 0.87) (Figure 6D). Collectively, these ROC analyses indicate that both gene- and miRNA-based candidates in fusion-positive and fusion-negative NSCLC display robust discriminatory power and may serve as informative biomarkers for subtype-specific diagnosis and patient stratification.

### CMap analysis identifies candidate compounds with therapeutic potential in fusion-positive and fusion-negative NSCLC

To identify small molecules with the potential to reverse subtype-specific gene expression signatures, we conducted a Connectivity Map (CMap) analysis using DEGs derived from fusion-positive and fusion-negative NSCLC samples. Candidate compounds were prioritized based on normalized connectivity scores (ncs), with negative scores indicating predicted antagonism of the disease-associated transcriptional profiles. In fusion-positive NSCLC, four compounds—MLN-4924, anisomycin, bosutinib, and AG-370—exhibited strong negative connectivity scores, suggesting potential capacity to counteract the fusion-positive gene expression program. MLN-4924, an inhibitor of NEDD8-activating enzyme (NAE), and anisomycin, targeting RPL37, were predicted to act on genes upregulated in fusion-positive tumors. Bosutinib targets CAMK1D and FRK, both of which were also upregulated in this subtype. Notably, AG-370 is a PDGFR inhibitor, consistent with our observation of significant dysregulation of PDGFR signaling in fusion-positive NSCLC (Table 5).

**Table 5.** List of drugs obtained from CMap Analysis based on normalised connectivity score (norm_cs), mechanism of action (MOA) and drug targets. Drugs* - selected based on the upregulated targets.

| Pert_iname | MOA | Targeted_upregulated_DEGs | Norm_cs |
| --- | --- | --- | --- |
| **Fusion-Positive** |  |  |  |
| MLN-4924* | Nedd activating enzyme inhibitor | NAE1 | -2.3 |
| Anisomycin* | DNA inhibitor | RPL37 | -2.05 |
| Bosutinib* | Src inhibitor, Abl inhibitor, Bcr-Abl inhibitor | CAMK1D, FRK | -2.02 |
| AG-370 | PDGFR inhibitor | — | -1.96 |
| **Fusion-Negative** |  |  |  |
| BIX-02189 | MEK inhibitor | — | -2.41 |
| Palbociclib | CDK inhibitor | — | -2.41 |
| AMG-232 | MDM inhibitor | — | -2.4 |
| Ro-4987655 | MEK inhibitor | — | -2.4 |
| PD-184352 | MEK inhibitor | — | -2.39 |
| RG-7388 | MDM inhibitor | — | -2.39 |
| Sulbutiamine | Acetylcholine receptor antagonist | — | -2.03 |
| BRD-A04553218* | Histamine receptor antagonist | SLC6A3 | -1.97 |

In fusion-negative NSCLC, eight compounds demonstrated strong inverse connectivity with the DEG signature, including BIX-02189, palbociclib, AMG-232, Ro-4987655, PD-184352, RG-7388, sulbutiamine, and BRD-A04553218. BIX-02189, Ro-4987655, and PD-184352 are MEK inhibitors, whereas AMG-232 and RG-7388 disrupt the MDM2–p53 interaction. Palbociclib functions as a CDK inhibitor, and sulbutiamine acts as an acetylcholine receptor agonist. These pathways were independently identified as dysregulated in fusion-negative tumors, supporting the relevance of the inferred compounds. BRD-A04553218 targets SLC6A3, a gene upregulated in fusion-negative NSCLC, further supporting its potential therapeutic relevance (Table 5). Overall, this analysis highlights subtype-specific small molecules with predicted capacity to reverse oncogenic transcriptional programs in fusion-positive and fusion-negative NSCLC, providing a rational basis for future experimental validation and therapeutic exploration.

## Discussion

We identified molecular features that distinguish EML4–ALK fusion-positive NSCLC tumors from fusion-negative counterparts, yielding insights into the biology of this oncogenic subtype and its therapeutic implications. Fusion-positive NSCLC was characterized by fusion-exclusive upregulated genes reflecting enhanced metabolic, secretory, and signal-transduction activity, with functional clusters enriched for intracellular signaling, cell proliferation, and glycolytic metabolism. The observed enrichment of glycolysis- and gluconeogenesis is consistent with the metabolic reprogramming commonly associated with aggressive tumor phenotypes ^59,60^.

Fusion-specific activation of the N-glycan biosynthetic pathway is of particular mechanistic interest, as elevated N-glycan expression correlates with metastatic potential across multiple tumor types ^61,62^. Dysregulated N-glycan biosynthesis has also been shown to potentiate cytokine signaling and tumor-promoting stress responses in diverse solid malignancies ^63–66^. We identified seven upregulated genes including TMEM258, P4HB, SEC24D, PDIA6, SIL1, DDOST, and HSP90B1 associated with ER protein processing in fusion-positive NSCLC. Among these, DDOST and TMEM258 were co-enriched within the N-glycan biosynthetic pathway, suggesting functional roles in metastatic behavior and therapeutic resistance specific to the fusion-positive subgroup. In parallel, upregulation of PKM, ALDOC, ENO2, and ALDH7A1, together with DPAGT1, a key regulator of N-glycan synthesis, supports the presence of a fusion-driven metabolic program that couples enhanced glycolysis with increased glycoprotein processing capacity. Such metabolic redirection may not only sustain rapid proliferation but also promote lactate accumulation and microenvironmental acidification, conditions known to facilitate tumor cell migration and invasion ^67^. Complementing these findings, elevated expression of GAMT, CHDH, and ALDH7A1 enzymes involved in glycine, serine, and threonine metabolism further reinforces the notion of broad metabolic plasticity underlying tumor adaptability and aggressiveness in fusion-positive NSCLC.

Beyond these metabolic alterations, GSEA confirmed significant upregulation of the amino sugar and nucleotide sugar metabolism pathway in fusion-positive tumors relative to fusion-negative cases. Leading-edge analysis identified six key upregulated genes including CYB5R1, FCSK, GFPT1, GMPPA, GMPPB, and GPI as principal contributors to this enrichment. Among these, GMPPA emerged as a particularly prominent node within the protein–protein interaction network. Together with DPAGT1 and DDOST, both of which also function as network hubs, GMPPA integrates multiple metabolic pathways, including glycolysis, N-glycan biosynthesis, and amino sugar metabolism. The convergence of these pathways highlights their potential utility as discriminative molecular features of fusion-positive NSCLC and positions them as candidate biomarkers or therapeutic targets. In addition, enrichment of the protein export pathway, marked by upregulation of SEC61A1, another PPI hub points to a heightened secretory phenotype associated with fusion-positive cases.

Furthermore, analysis of downregulated DEGs specific to fusion-positive tumors revealed marked suppression of signaling and adhesion-related pathways, including calcium signaling and extracellular matrix (ECM)–receptor interaction, both of which are central to cellular communication, polarity, and migration. Key hub genes within these networks, including COL4A2, PDGFRB, and LAMA4, were strongly repressed. Calcium signaling plays a fundamental role in regulating cytoskeletal dynamics, cell motility, and adhesion turnover, and spatial calcium gradients are known to coordinate directional migration ^68–70^. Likewise, ECM remodeling is essential for tumor invasion and metastasis, providing both structural support and biochemical cues that shape tumor–stroma interactions ^71–74^. The observed downregulation of ECM–receptor and focal adhesion components therefore suggests impaired cell–cell and cell–matrix adhesion, a state that may paradoxically facilitate cellular detachment and dissemination. This suppression of adhesion-associated programs occurs alongside upregulation of N-glycan biosynthetic pathways, collectively implying that altered glycosylation may disrupt normal adhesion dynamics while promoting metastatic competence ^75,76^. Concurrent attenuation of calcium influx, a key regulator of adhesion assembly and turnover, may further reinforce this effect, establishing a microenvironment permissive to invasion and dissemination ^77^.

Beyond adhesion and cytoskeletal regulation, several immune- and cytokine-associated signaling pathways were also suppressed in fusion-positive NSCLC. Negatively enriched pathways were observed in fusion-positive versus fusion-negative tumors including cytokine-receptor, ECM-receptor, calcium signaling, cancer pathways, and Toll-like receptor signaling, with PDGFRB emerging as a shared node across these modules. The cytokine–cytokine receptor network plays an important role in modulating tumor aggressiveness and shaping immune responsiveness, and its attenuation has been linked to impaired immune activation and enhanced metastatic potential ^78,79^. Accordingly, downregulation of PDGFRB and associated signaling components in our dataset may reflect a compromised cytokine milieu that favors immune evasion in ALK fusion–driven tumors. Several downregulated hub genes including PDGFRB, COL4A2, LAMA4, and CTSK converged across multiple functional pathways, encompassing cytokine–cytokine receptor interaction, ECM–receptor interaction, gap junction communication, focal adhesion, Toll-like receptor signaling, and calcium signaling (Figure 2D). This convergence highlights a broad suppression of intercellular communication and matrix sensing, processes that are critical for maintaining tissue integrity and immune surveillance. Such coordinated downregulation may confer a survival advantage under stress conditions and facilitate metastatic dissemination.

Furthermore, we integrated fusion-gene expression data with microRNA profiles to delineate fusion-associated microRNA dysregulation. hsa-miR-3065 emerged as exclusively upregulated in fusion-positive tumors and was predicted to target six hub genes: PDGFRB, CTSK, COL4A2, SPARC, FBN1, and LUM (Figure 3D). While altered expression of hsa-miR-3065 has been reported in esophageal squamous cell carcinoma, renal cell carcinoma and aggressive early-stage lung adenocarcinoma ^80–83^, its functional role in in EML4–ALK-driven NSCLC has remained largely unexplored.

COL4A2 and PDGFRB, predicted target of hsa-miR-3065 are components of calcium signaling, ECM–receptor interaction, and cytokine–cytokine receptor interaction pathways, all of which are central to tumor cell motility, adhesion, and immune modulation. Disruption of calcium homeostasis is a well-established feature of cancer progression and metastasis ^84^. PDGFRB has been implicated in invasive and metastatic behavior in colorectal and hepatocellular carcinomas ^85,86^, whereas COL4A2 has been described as an inhibitor of angiogenesis and tumor progression in lung cancer ^87^. From a therapeutic perspective, modulation of calcium signaling has been shown to constrain tumor proliferation, metastasis, and angiogenesis, while potentially enhancing anti-tumor immune responses ^88–90^. Another predicted hsa-miR-3065 target, CTSK, intersects with Toll-like receptor (TLR) signaling and has been implicated in tumor-promoting processes through extracellular matrix degradation and angiogenesis in multiple malignancies ^91,92^. SPARC modulates cell adhesion and proliferation through multiple signaling pathways, and reduced SPARC expression has been associated with poor prognosis in lung cancer ^93^. FBN1 contributes to extracellular matrix architecture and has been linked to structural remodeling in cancer-associated stroma ^94^. LUM, a stromal ECM proteoglycan, correlates with cancer-associated fibroblast abundance and immune-evasive stromal states, consistent with our observed CAF-infiltration patterns in fusion-positive NSCLC ^95^. Structural modeling with AlphaFold 3 reliably placed hsa-miR-3065-3p on the COL4A2, FBN1, PDGFRB, and LUM 3′-UTR in an AGO2-compatible configuration, supporting the predicted interaction. ROC analysis further showed that hsa-miR-3065, together with PDGFRB, COL4A2, SPARC, and LUM, achieved AUC values >0.7, supporting their potential to distinguish fusion-positive tumors.

In the fusion-negative NSCLC cohort, we identified seven distinct upregulated differentially expressed miRNAs including hsa-miR-21-5p, hsa-miR-141-3p, hsa-miR-431-5p, hsa-miR-199b-5p, hsa-miR-148a-3p, hsa-miR-323b-5p, and hsa-miR-889-3p that collectively targeted a set of downregulated hub genes, including MAP3K19, DNAAF6, DTHD1, ZBBX, TEKT1, and TEKT2. In parallel, six downregulated miRNAs—hsa-let-7a-2-3p, hsa-let-7d-3p, hsa-let-7d-5p, hsa-let-7e-5p, hsa-let-7f-1-3p, and hsa-let-7f-2-3p—were predicted to regulate seven upregulated hub genes: CDC25C, TRIP13, MELK, KIF23, TTK, PBK, and NCAPG2. Among these, CDC25C, a key G2/M checkpoint phosphatase, has been associated with unfavorable outcomes in NSCLC ^96,97^. TRIP13 promotes tumor progression through activation of AKT, mTORC1, and c-Myc signaling ^98–100^ and together with KIF23 has been proposed as a therapeutic target linked to aggressive disease behavior and poor prognosis ^101,102^. Overexpression of TTK has similarly been correlated with reduced overall survival ^103,104^. Consistent with these observations, an eight-miRNA upregulated signature—hsa-miR-21-5p, hsa-miR-141-3p, hsa-miR-431-5p, hsa-miR-199b-5p, hsa-miR-148a-3p, hsa-miR-323b-5p, hsa-miR-889-3p, and hsa-miR-592—combined with downregulation of three let-7 family members (hsa-let-7a-2, hsa-let-7f-1, and hsa-let-7f-2) demonstrated reliable diagnostic performance in ROC analyses and may serve as candidate biomarkers for fusion-negative NSCLC. These findings are concordant with previously reported miRNA biomarkers in lung adenocarcinoma, including miR-10b, miR-126, miR-130a, miR-505, and miR-4529 ^105–108^.

The upregulated miRNA identified here, notably hsa-miR-21-5p has been implicated in NSCLC progression and is under consideration for clinical targeting ^109^. hsa-miR-141-3p overexpression has been reported to inhibit cell proliferation, migration and invasion of NSCLC cells ^110^. Other miRNAs displayed context-dependent roles. hsa-miR-199b-5p has been reported to act as an oncogenic in gastric cancer ^111^ while exhibiting tumor-suppressive functions in triple-negative breast cancer ^112^ and acute myeloid leukemia ^113^. hsa-miR-148a-3p has been linked to tumorigenesis in prostate, glioblastoma and osteosarcoma ^114,115^, and may inhibit epithelial–mesenchymal transition in NSCLC depending on cellular context ^116^. hsa-miR-889-3p promotes proliferation through immune-gene suppression in osteosarcoma ^117^. In contrast, hsa-miR-323b-5p and hsa-miR-889-3p remain relatively understudied in NSCLC, underscoring the need for targeted functional validation. Finally, the let-7 family, a canonical tumor-suppressive miRNA axis in lung cancer was consistently downregulated in the fusion-negative cohort, in agreement with reports linking let-7 loss to shortened survival and poor prognosis in NSCLC ^118,119^.

To directly interrogate fusion-linked regulatory mechanisms, we also examined miRNAs predicted to target the parental EML4 and ALK transcripts. Because fusion events disrupt the native EML4 3′-UTR, miRNAs that normally bind EML4 may lose their primary binding site and become redistributed toward alternative targets. Consistent with this model, we identified five EML4-targeting miRNAs—hsa-miR-29c-3p, hsa-miR-29a-3p, hsa-miR-152-3p, hsa-miR-199b-3p, and hsa-miR-627-5p that were also predicted to regulate six hub genes in fusion-positive NSCLC (COL4A2, FBN1, NID2, PDGFRB, SPARC, and LAMA4). In parallel, the ALK-targeting miRNA hsa-miR-6501-3p was predicted to regulate HYOU1, an upregulated hub gene in fusion-positive tumors. miR-29c-3p has previously been proposed as diagnostic biomarkers for lung cancer^120^. hsa-miR-199b-3p has been associated with poor prognosis and immune suppression in gastric cancer while exhibiting tumor-suppressive activity in osteosarcoma ^121^. Although hsa-miR-6501-3p has been correlated with NSCLC prognosis, its functional role remains incompletely defined and warrants investigation ^122^.

In parallel with transcriptomic and miRNA analyses, we examined the tumor microenvironment to assess how EML4–ALK fusions influence stromal and immune organization in NSCLC. TME-mediated resistance to ALK inhibition has emerged as a clinically relevant factor in EML4–ALK-positive cases, with stromal and immune interactions modulating the efficacy of tyrosine kinase inhibitors ^123,124^. Cancer-associated fibroblasts (CAFs) are central contributors to immune evasion, angiogenesis, extracellular matrix remodeling, and metastatic dissemination processes that collectively undermine therapeutic durability ^125,126^. Our analyses revealed reduced CAF infiltration in fusion-positive tumors relative to fusion-negative NSCLC, consistent with an attenuated stromal program in the fusion-positive subset. Correlation analyses supported this pattern (Figure 5). Seven downregulated genes—NID2, PDGFRB, FBN1, COL4A2, LAMA4, LUM, and AEBP1—were positively correlated with CAF abundance, whereas upregulated HYOU1 showed negative correlations with CAF infiltration in fusion-positive tumors. These relationships are concordant with the observed downregulation of ECM and adhesion-related circuits and suggest that EML4–ALK-driven tumors may rely less on desmoplastic stromal scaffolding and more on intrinsic metabolic and secretory reprogramming. In keeping with classical CAF biology characterized by sustained activation, ECM deposition, and stromal desmoplasia—our transcriptomic analyses showed that genes downregulated in fusion-positive NSCLC converge on downregulated receptor interaction activity, focal adhesion, calcium signaling and actin cytoskeleton regulation, pathways integral to CAF function and matrix remodeling (Figure 2). Previous studies have shown ECM-associated signatures to stromal compartments, including CAFs, reinforcing their contribution to tumor behavior ^127–129^. Functionally, the downregulated hub genes, most notably NID2, PDGFRB, FBN1, and COL4A2 map to extracellular matrix structure and cell adhesion programs that are critical for tumor migration and invasion. These nodes intersect with epithelial–mesenchymal transition (EMT) processes, which are characterized by reduced epithelial adhesion and increased motility ^130^. Clinically, diminished expression of adhesion-related genes has been associated with distant metastasis in lung cancer, and reduced expression of LAMA family members, including LAMA4, has been linked to poor overall survival in cancers ^131,132^.

To translate these regulatory networks into testable hypotheses, we performed Connectivity Map (CMap) analyses to identify small molecules predicted to counteract fusion-positive transcriptional dysregulation. This analysis highlighted several candidate compounds, including MLN-4924 (pevonedistat), anisomycin, bosutinib, and AG-370. MLN-4924, an inhibitor of protein neddylation, is currently under clinical evaluation in both solid and hematologic malignancies ^133,134^. Anisomycin, originally characterized as an inhibitor of protein and DNA synthesis ^135^, has been reported to sensitize NSCLC cells to chemotherapy and EGFR-targeted agents through inhibition of the PI3K/Akt/mTOR pathway ^136^. AG-370, a tyrosine kinase inhibitor targeting PDGF signaling, suppresses PDGF-driven autophosphorylation and fibroblast mitogenesis, a signaling axis that emerged as dysregulated in our fusion-positive dataset ^137^. Together, these compounds converge on pathways related to ER stress and protein handling, proliferative signaling, and stromal PDGF circuitry, collectively nominating metabolic–secretory–stromal interactions as a potentially tractable vulnerability in EML4–ALK fusion-positive NSCLC.

In the fusion-negative cohort, CMap analysis implicated BIX-02189, a MEK5/ERK5 inhibitor, palbociclib, a CDK4/6 inhibitor, as well as AMG-232, Ro-4987655, PD-184352, RG-7388, sulbutiamine, and BRD-A04553218 as candidate therapeutic leads. Notably, BIX-02189 and palbociclib have demonstrated synergistic anti-proliferative effects with temozolomide in glioma models through suppression of GINS2 and DNA damage response pathways ^26,138^. AMG-232 and Ro-4987655 are under investigation in acute myeloid leukemia and solid tumors, respectively ^139,140^, while PD-184352 (CI-1040) has been evaluated across multiple cancer types, including lung, breast, pancreatic, and colorectal malignancies ^141,142^. RG-7388 (idasanutlin), an MDM2 antagonist, is also undergoing clinical assessment in both hematologic and solid tumors ^34,143^. Sulbutiamine, a thiamine derivative, is clinically used for the management of asthenia ^144^. BRD-A04553218, an H1 histamine receptor antagonist commonly employed for allergic conditions ^145^, was notable for targeting SLC6A3, a gene upregulated in our fusion-negative cohort, suggesting a noncanonical and context-dependent avenue for therapeutic modulation that warrants further mechanistic investigation.

We acknowledge that the relatively small number of fusion-positive cases limits statistical power and may constrain generalizability. All regulatory inferences presented here including miRNA–mRNA targeting relationships and structural compatibility assessments are computational in nature and will require experimental validation. In addition, reliance on bulk transcriptomic data restricts resolution of cell-type–specific effects within the tumor microenvironment. Future studies incorporating functional perturbation approaches, spatial and single-cell profiling, and prospective clinical validation will be necessary to substantiate the mechanistic roles of the identified miRNA–mRNA circuits and to evaluate their translational relevance.

## Conclusion

This study delineates distinct molecular and immunological landscapes between EML4-ALK fusion-positive and fusion-negative lung adenocarcinoma. Fusion-positive tumors exhibit metabolic reprogramming, reduced stromal infiltration, and miRNA-mediated repression of ECM-related genes, particularly via *hsa-miR-3065*. In contrast, fusion-negative tumors are characterized by proliferative signatures and let-7 family downregulation. We identified a distinct set of fusion-positive exclusive upregulated hub genes, including HYOU1, DPAGT1, and DDOST, which play critical roles in glycolysis, N-glycan biosynthesis, and protein export. Fusion-specific downregulated genes such as NID2, SPARC, PDGFRB, FBN1, COL4A2, LAMA4, LUM, and AEBPT1 were associated with pathways including focal adhesion, calcium signaling, ECM receptor interaction, cytokine-cytokine receptor interaction, and TLR signaling. Notably, hsa-miR-3065 was upregulated in fusion-positive NSCLC and was found to regulate six key hub targets including PDGFRB, CTSK, COL4A2, SPARC, FBN1, and LUM, all of which are integral to dysregulated pathways. ROC analyses demonstrated that hsa-miR-3065 and its targets, including HYOU1, LAMA4, SPARC, PDGFRB, COL4A2, and LUM, possess effective predictive power, making them promising prognostic biomarkers for fusion-positive NSCLC. Additionally, reduced infiltration of CAFs in fusion-positive NSCLC was associated with positive correlations to NID2, SPARC, PDGFRB, FBN1, COL4A2, LAMA4, LUM, and AEBP1, and negative correlation with HYOU1. These findings highlight the distinct tumor microenvironment architecture of fusion-positive NSCLC and support the hypothesis that such differences may underline resistance to targeted therapies. Overall, the identified genes and miRNAs provide a valuable molecular framework for the development of diagnostic markers and targeted therapies, offering potential for personalized treatment strategies and improved clinical outcomes in fusion-positive NSCLC.

## CRediT Authorship Contribution Statement

Divya Mishra: Methodology, Software, Data curation, Formal analysis, Visualization, Investigation, Writing - Original draft, review & editing. Shivangi Agrawal: Software, Formal analysis, Investigation, Visualization, Writing - Original draft. Divya Malik: Software, Formal analysis, Investigation, Writing - Original draft. Ekta Pathak: Conceptualization, Formal analysis, Investigation, Visualization, Writing - original draft, Writing - review & editing. Rajeev Mishra: Conceptualization, Methodology, Investigation, Writing - original draft, Writing - review & editing, Visualization, Supervision, Project administration.

## Acknowledgments

We gratefully acknowledge the following sources of support: D.M. for the support from DBT-JRF and SRF; S.A. for project assistant support from DST-CURIE-2022-80, Government of India; Divya for CSIR-JRF and SRF from Govt. of India. R.M. gratefully acknowledges for the research support from DST-CURIE-2022-80(G), Government of India. We acknowledge BioRender.com, whose tools were used for enhancing the clarity of some figures.

